# Rapid, in-patient adaptations of *Legionella pneumophila* to the human host

**DOI:** 10.1101/2022.05.23.492981

**Authors:** Daniël Leenheer, Anaísa B. Moreno, Susan Murray, Sophie Jarraud, Christophe Ginevra, Lionel Guy

## Abstract

*Legionella pneumophila* are host-adapted bacteria that infect and reproduce primarily in amoeboid protists. Using similar infection mechanisms, they infect human macrophages, and cause Legionnaires’ disease, an atypical pneumonia, and the milder Pontiac fever. We hypothesized that, despite these similarities, the hosts are different enough so that there exist high-selective value mutations that would dramatically increase the fitness of *Legionella* inside the human host. By comparing a large number of isolates from independent infections, we identified two genes, mutated in three unrelated patients, despite the short duration of the incubation period (2-14 days). One is a gene coding for an outer membrane protein (OMP) belonging to the OmpP1/FadL family. The clinical strain, carrying the mutated OMP homolog, grows faster in macrophages than the wild type strain, and thus appears to be better adapted to the human host. The other is a gene coding for a protein involved in cyclic-di-GMP regulation, which in turn modulates flagellar activity. As human-to-human transmission is very rare, fixation of these mutations into the population and spread into the environment is unlikely. Therefore, convergent evolution – here mutations in the same genes observed in independent human infections – could point to adaptations to the accidental human host. These results suggest that despite its ability to infect, replicate, and disperse from amoebae, *L. pneumophila* is not well adapted to the human host.

**Impact statement:** *Legionella pneumophila* is primarily infecting amoeboid protists, but occasionally infects human lung macrophages, causing Legionnaires’ disease, an atypical pneumonia. By comparing 171 isolates from patients to their probable environmental source, we identified 119 mutations that presumably occurred in-patient. Among these, several mutations occurred in the gene. In particular, two genes were mutated thrice, significantly more often than expected by chance alone, and are likely to represent adaptations to the human host. We experimentally show that, for one mutation at least, the mutated strain grows faster in human macrophages than in amoebae. By specifically investigating in-patient mutations, we were able to identify two genes that might be involved in human host-specific adaptations of *L. pneumophila*. This result suggests that *L. pneumophila* is not particularly well adapted to the human host, as mutations get fixed in-patient, during the short course of an infection (2-14 days), indicating a very high selective value.

**Data Summary:** The sequencing data generated in this study are available in the NCBI database under the BioProject accession number: PRJEB52976.

## Introduction

### Legionella

*Legionella* are Gram-negative bacteria, belonging to the Gammaproteobacteria, commonly found in aquatic and soil environments ([1] where they are able to infect a wide variety of protozoan hosts ranging from free-living amoebae to ciliated protozoa [2, 3]. Inside these hosts, *Legionella* is able to resist killing by water disinfection procedures commonly employed in man-made potable water systems [4]. Human exposure commonly occurs via the inhalation of contaminated aerosols produced from these systems, through showers, taps, fountains, etc. In the past two decades, cases of Legionnaires’ disease have dramatically increased, with an estimated 8-10 fold surge between 2000 and 2018 [5]. The fatality rate is high, about 8%, with the elderly, males, smokers, and the immunosuppressed are at higher risks to contract the disease.

Several *Legionella* species are known to be capable of infecting mammalian cells, such as alveolar macrophages inside the human lung, with *L. pneumophila* being the most frequent human pathogen. In susceptible cases, *Legionella* infection may lead to a severe, atypical pneumonia known as legionellosis or Legionnaires’ disease, or the milder Pontiac fever [6, 7]. Legionnaires’ disease typically lasts 2 to 14 days, and ends up by *Legionella* being cleared by the immune system, or by the death of the patient [8]. Except for one documented case, human-to-human transmission appears to be very rare [9].

### Drivers of genomic diversity

Estimates of bacterial substitution rates were initially believed to be in the order of 10^-10^ to 10^-9^ substitutions per site per year [10, 11]. However, studies have shown that short-term evolution rates are more likely to be in the order of 10^-9^ to 10^-5^ substitutions per site per year [12, 13]. With the advances in whole genome sequencing, it has become possible to sequence multiple isolates from the same host. This has led to several studies comparing pairs of genomes from within the same host, such as *Mycobacterium tuberculosis* [14], *Escherichia coli* [15], *Clostridium difficile* [16], *Staphylococcus aureus* [17], *Klebsiella pneumoniae* [18], and *Helicobacter pylori* [19], with within-host point mutation rates ranging from 0.5 to 30 mutations per genome per year. The evolutionary rate of *L. pneumophila* ST578 strains from Alcoy is estimated at ~10^-5^ substitutions per site per year [20]. This translates to a lower bound estimate of approximately 35 single nucleotide polymorphisms (SNPs) per genome per year, or 0.2-1.3 SNP per genome for the average (2-14) day incubation period. These rates are however for substitutions, with mutation rates (and especially high-selective value mutations) probably occurring at higher rates.

### Recombinations in *L. pneumophila*

Substitutions occur through point mutation, but also through recombination, following transduction, conjugation and transformation. All three mechanisms have been described in *L. pneumophila* [21, 22], and comparative genomics revealed that recombination is actually responsible for most substitutions in this organism [23–25]. The majority of recombinations occurs in a few hot-spots, which include regions encoding outer membrane proteins, the lipopolysaccharide (LPS) region and effectors secreted by the Type IVB Secretion System (T4BSS) [24].

### Adaptations to the human host

This study aims to identify human-specific adaptations to the human host in *L. pneumophila*. While the mechanisms used to infect amoebae and human alveolar macrophages are similar, differences exist. What these differences are, what role they play, and how important they are, is currently not well known [26, 27]. Since human-to-human transmission appears to be a very rare event, with only a single case documented thus far [9], any human-specific adaptations are unlikely to be fixed and spread, either to other human hosts or to the environment. Therefore, we hypothesize that some mutations, when arising during the short incubation period (2-14 days), would give a strong selective advantage in infecting and colonizing the human host. Substitutions that proved beneficial in various amoebae have very different effects on the fitness of *Legionella* in human cells [28], and it is difficult to predict whether specific substitutions will be beneficial to *Legionella* when infecting the human host. However, substitutions which would reduce the *Legionella*’s probability of being recognized by the immune system are likely to be favored in human infections, and to have a large selective value.

To identify these mutations and separate them from neutral mutations we compared 171 pairs of strains, one from a single infection or an outbreak, the other from its inferred environmental source. By comparing these clinical samples to their respective environmental samples, we identified adaptations occurring in the same gene in independent infections.

## Results

### Isolate pairing

We sequenced the genomes of 166 clinical and environmental samples belonging to *L. pneumophila* using the Illumina MiSeq platform. These strains were isolated from clinical and environmental sources sampled during the investigation of sporadic cases in Sweden and France, as well as from outbreaks in Madrid (1996) [29] and Murcia (2001) [30], Spain. Most strains corresponded to serogroup 1, but sequence types varied (**Supplementary Table 1**). Additional samples, including genomic data, were retrieved from published outbreak investigations [20, 24, 31–36]. Each clinical sample was paired with its closest environmental relative, to form a comparison. Comparisons potentially caused by co-infections or independent infections within a short time period were removed from our final analysis [31, 37, 38]. After filtering, we obtained 171 comparisons (**Table 1**, **Supplementary Table 2**), of which 100 come from isolates sequenced in this study, and 71 from publicly available data. In total, 24 comparisons come from 2 separate outbreaks and 147 from isolated Legionnaires’ disease cases.

**Table 1:** Summary of the comparisons between environmental and clinical strains analyzed in this study.

| Sample origin | Comparisons |
| --- | --- |
| Madrid 1996 outbreak | 7 |
| Murcia 2001 outbreak | 17 |
| Folkhälsomyndigheten | 18 |
| National Legionella Reference Centre, Lyon | 58 |
| Published | 71 |
| **Total** | **171** |

A phylogeny based on the core genome shared by 95% of all samples confirmed the similarity between the pairs of samples and among the isolates from outbreaks (**Supplementary Figure 1**). The supplementation of the phylogeny with strains Alcoy, Corby, Lens, Lorraine, Paris, and Philadelphia reveal that many strains are closely related to *L. pneumophila* Paris.

**Supplementary Figure 1:**
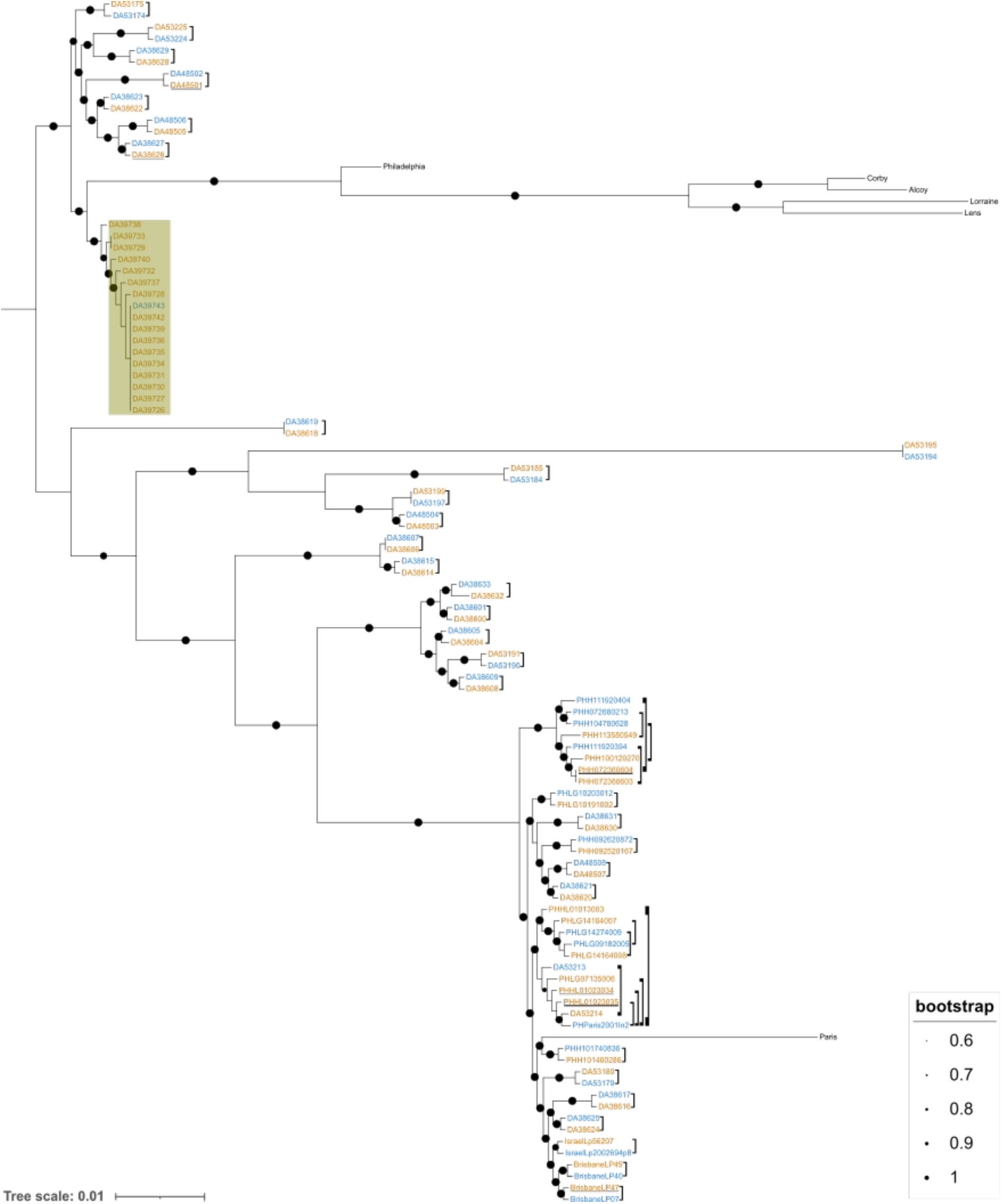
Unrooted maximum-likelihood phylogenetic tree of *Legionella pneumophila*. All samples in this study, as well as a few reference ones (strains Philadelphia, Lorraine, Lens, Corby, and Alcoy) are included. Environmental isolates are shown in blue, clinical isolates in orange. Comparisons from single cases are shown with brackets, and outbreaks are highlighted in olive. Samples underlined in black contain mutations in an EAL domain containing protein, samples underlined in grey contain mutations in a membrane protein. The tree is based on a Legionella-specific cgMLST scheme, resulting in an alignment of 2 253 410 nucleotide positions. Circles on branches represent the percentage of bootstrap trees supporting the node. Bootstrap support values under 60 are not shown. The scale represents the average number of substitutions per site.

### Genomic variation between pairs of isolates

To investigate what genetic variants might have occurred in-patient, reads from environmental isolates were assembled de novo and reads from the corresponding clinical strain were mapped to the assembled environmental isolate. Recombining regions were removed from the analysis. Comparisons considered in this study had at most 20 substitutions, and in 16 comparisons, no differences could be found. In total, 206 substitutions (173 SNPs and 33 short indels) were identified in the 171 comparisons. Among indels, 2 (6%) were in-frame. Among the 173 SNPs, 123 (71%) were non-synonymous and the remaining 50 (29%) were synonymous; in 5 (3%) of the cases, the substitution resulted in a premature stop codon. Ninety (52%) were transitions and 83 (48%) were transversions.

### Simulations

To simulate the number of SNPs per gene that could be expected in an environment without selective pressure, 1000 studies were simulated, where 123 SNPs (total number of intragenic non-synonymous SNPs found in this study) were randomly distributed to the 3033 genes harbored by *L. pneumophila* strain Paris (**Figure 1**). Among the 1000 simulations, only 3 had 7 or 8 genes mutated twice, while none had 2 genes mutated 3 times. The probability of obtaining 7 or more genes mutated twice, respectively 2 genes mutated three times, from the simulated random distribution was estimated by one-sided Mann-Whitney *U* tests, using a single value as one of the samples. Seven genes or more mutated twice was significantly more than the distribution obtained by random sampling (*p* = 0.038), although only marginally. On the other hand, two genes mutated three times was significantly more than expected from the random distribution (*p* < 10^-6^). These results indicate that genes are likely not mutated randomly, but that there is an excess of genes mutated multiple times, suggesting that convergent evolution is at play.

**Figure 1:**
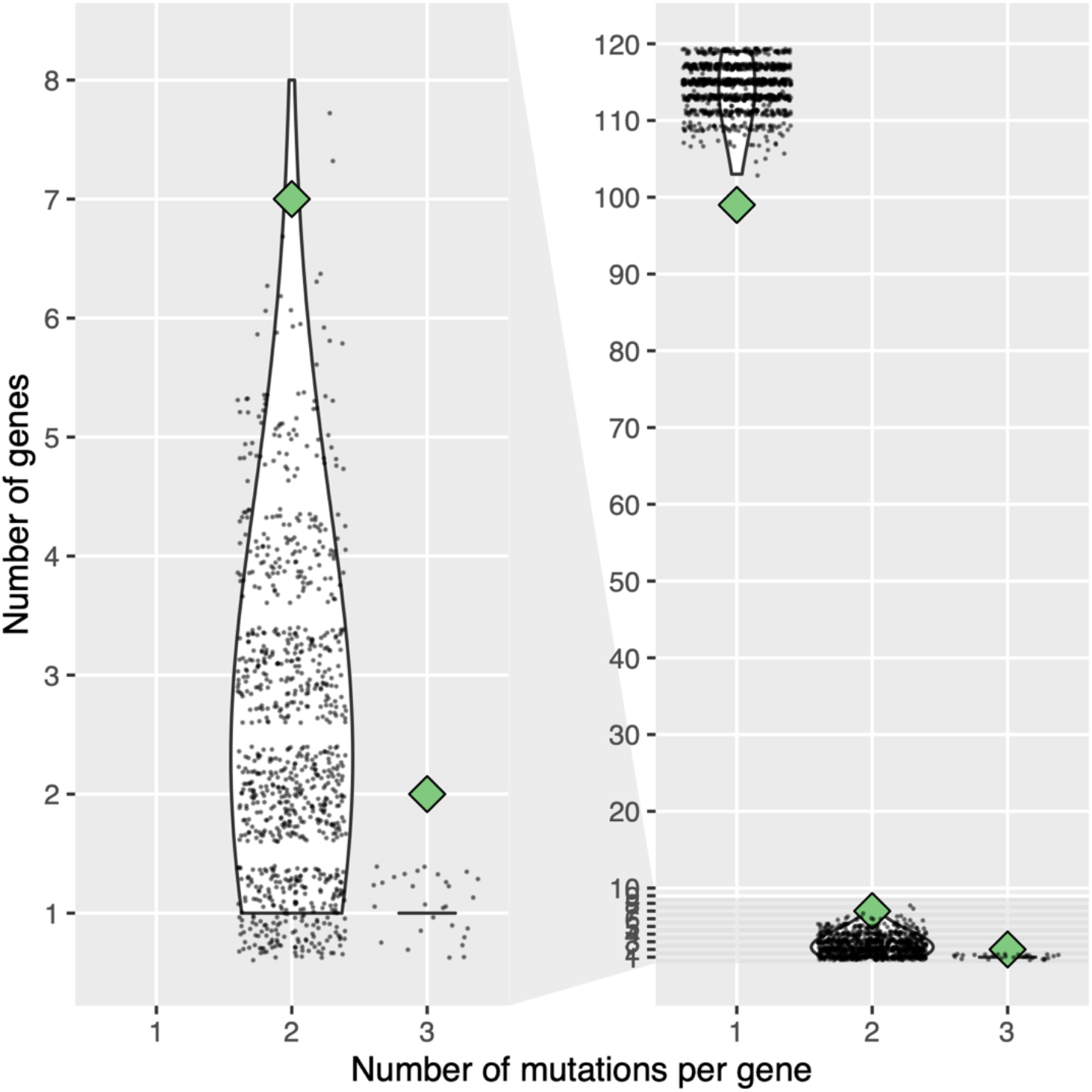
Number of mutations per gene in this study compared to 1000 simulations based on random sampling. The left panel is a subset of the right panel. The number of genes mutated once or more in comparisons between an environmental and clinical isolate observed in this study are displayed with green diamonds. The distribution of the number of genes mutated the same number of times in 1000 random sampling of 119 SNPs among the 3033 genes of *L. pneumophila* strain Paris is displayed with violin plots. Individual data points are overlaid as a cloud of points.

### Convergently mutated genes

To investigate which genes were mutated in multiple independent infections or outbreaks we linked our variants to orthologous gene families obtained with OrthoMCL. Variants occurring in identical families in two or more comparisons, and therefore likely to represent instances of convergent evolution can be seen in **Table 2**. Of particular interest are genes mutated independently in three separate comparisons: one encodes an (outer) membrane protein (lpg0707/lpp0762/lpl0744), and the other one for a EAL domain-containing protein (lpg0891/lpp0952/lpl0922). The outer membrane protein belongs to the OmpP1/FadL family. Two of the mutations led to proteins being 97 and 60% shorter than the wild-type protein by introducing a stop codon, while the third mutation did not lead to a change in secondary structure. The EAL-containing domain protein was also mutated three times, but none of the three mutations resulted in a shortened protein. These two genes are referred to as potentially adaptive in the human host (PAHH).

**Table 2:** Genes mutated more than once in clinical strains of *L. pneumophila*.

| Gene | Locus tag / Accession | Mutations in clinical isolates | Number of mutations |
| --- | --- | --- | --- |
| EAL domain-containing protein | lpg0891 | PHH072360604:G->R<br>PHHL01023035:G->D<br>DA48501:I->T | 3 |
| (outer) membrane protein | lpg0707 | BrisbaneLP47:G->*<br>PHHL01023034:W->*<br>DA38626:G->D | 3 |
| Efflux RNS transporter permease subunit | lpg1096 | IsraelLp56207:G->D<br>DA38604:F->F | 2 |
| KaiB-like protein 2 | WP_027224181.1 | DA38608:N->I<br>DA53225:INDEL | 2 |
| DUF1566 domain-containing protein | WP_014844754.1 | DA38600:W->*<br>DA38632:INDEL | 2 |
| GTP diphosphokinase | lpg2009 | DA53180:G->V<br>DA53214:G->V | 2 |
| Glycosyltransferase family 2 protein | lpg2478 | DA38604:R->S<br>DA38600:INDEL | 2 |
| ATP-dependent helicase/deoxyribonuclease subunit B | lpg2269 | PHLG14164007:S->N<br>PHLG14164008:S->N | 2 |
| CAAX amino terminal protease | lpg1525 | DA38604:INDEL<br>DA38608:INDEL | 2 |

### Growth assays

To determine the *in vitro* effect of the PAHH variants, two clinical strains (DA38626 and DA48501), two environmental strains (DA38627 and DA48502), together with *L. pneumophila* Paris (DA57510) as a reference strain were transformed with a plasmid encoding a YFP fluorescent protein. DA38626 carries a mutation in the OMP/FadL homolog, while DA48501 carries one in the EAL-containing protein. The corresponding environmental strains are otherwise isogenic to the clinical strains, although we cannot exclude that some structural variations were missed, due to the sequencing method used (Illumina). The plasmid does not markedly impact the growth rate of any of the strains (**Supplementary Figure 2**).

**Supplementary Figure 2:**
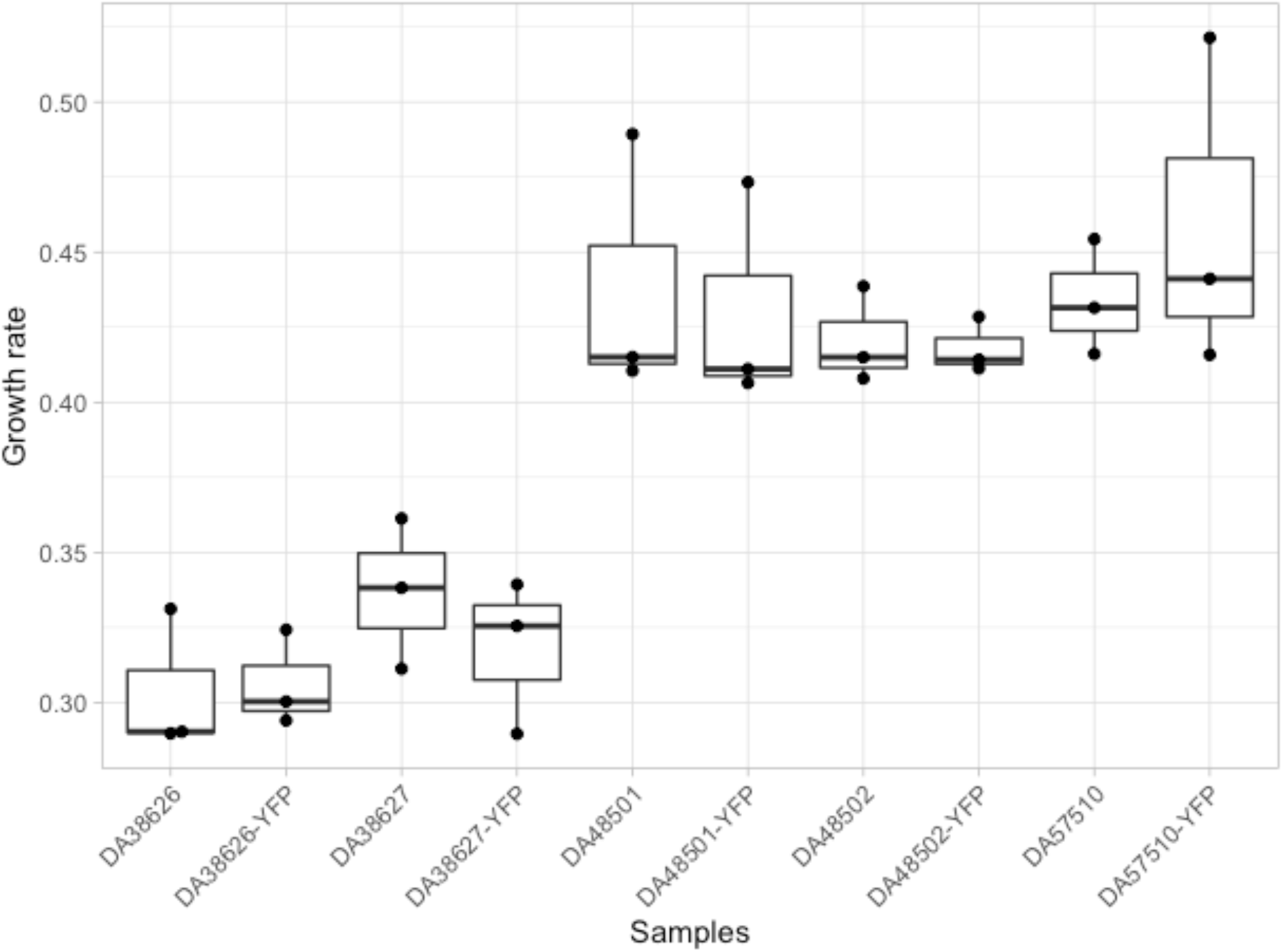
Comparison of growth rates of the five strains used in this study, with and without the fluorescence-carrying plasmid. Each dot represents the average of three technical replicates, and each strain was measured with three separate biological replicates. Boxplots provide a visual estimate of the distribution. DA38626 (clinical isolate) carries a mutated allele of the OmpP1/FadL homolog (lpg0707), while DA38627 (environmental) has the wild-type allele. DA48501 (clinical isolate) carries a mutated allele of the EAL-containing protein (lpg0891), while DA48502 has the wild-type one. DA57510 is *L. pneumophila* str. Paris.

To assess the effect of the PAHH mutations, the two pairs of isolates were first grown extracellularly, in liquid medium (**Supplementary Figure 3**). The pairs of clinical and environmental isolates exhibited very similar growth within the pair. The mutation in the OmpP1/FadL homolog in DA38626 resulted in a small decrease in growth rate (3.9% lower for DA38626, resulting in 3.4 minute increase in doubling time). The mutation in the EAL-containing protein resulted in a more important reduction of growth rate (16.7% lower growth rate, resulting in 15.5 minutes increase in doubling time. The reference Paris strain (DA57510) displayed an intermediate growth rate (0.412, doubling time of 101 minutes).

**Supplementary Figure 3:**
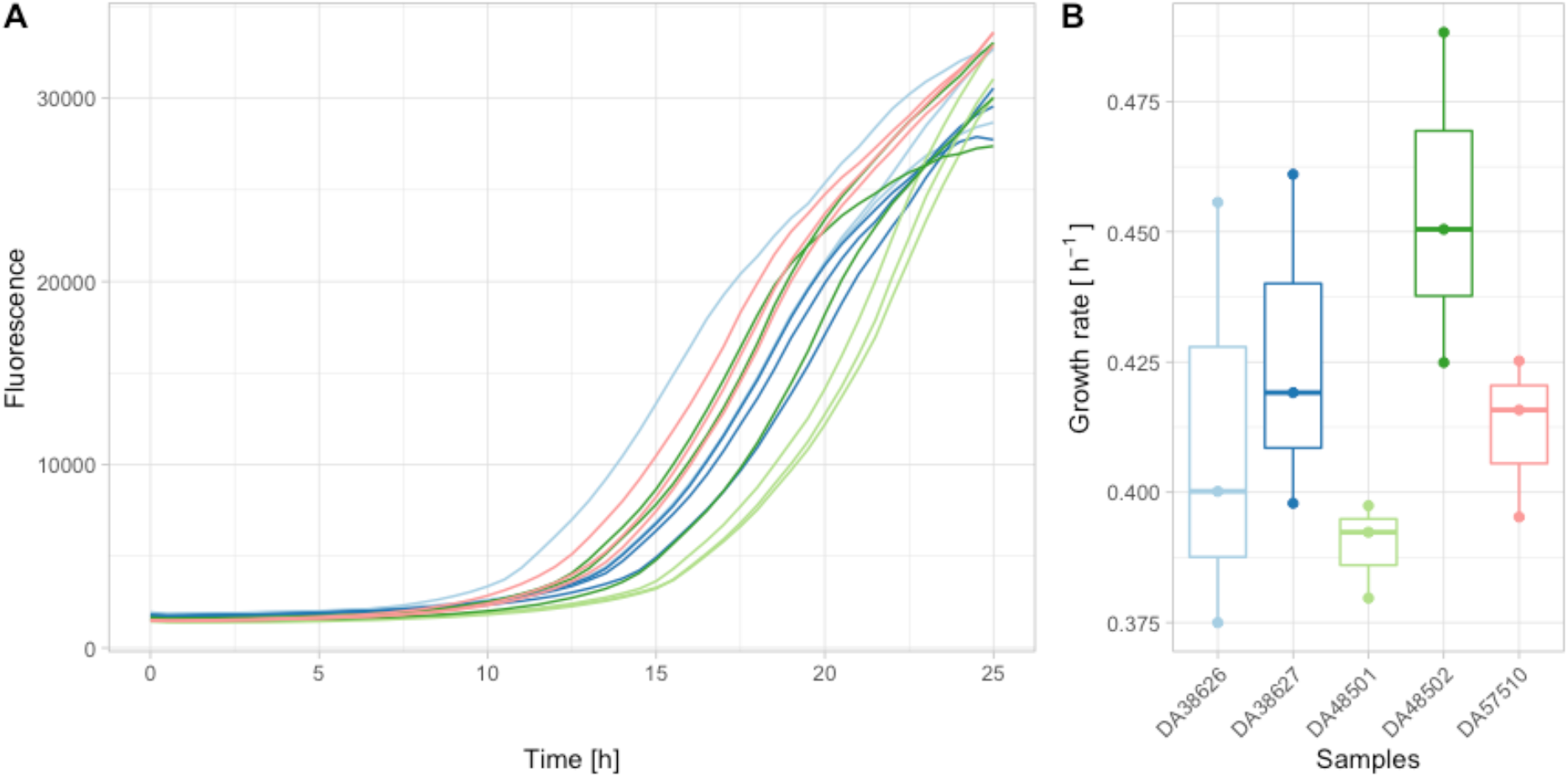
Growth of the two strains carrying a potential adaptation to the human host, compared to their wild types. **A:** Growth curves, as measured by the fluorescence emitted by the strains (y-axis). The x-axis is the time (in hours). Each curve is the average of three technical replicates. **B:** Boxplots of growth rates. Each dot represents the growth rate of the average growth curve shown in A, as calculated by the R package Growthcurver[39]. DA38626 (clinical isolate, pale blue) carries the mutated OmpP1/FadL homolog (lpg0707), while DA38627 (environmental, dark blue) has the wild type gene. DA48501 (clinical isolate, pale green) carries a mutated EAL-containing protein (lpg0891), while DA48502 (environmental, dark green) has the wild type gene. DA57510 (red) is *L. pneumophila* str. Paris and serves as control.

The intracellular replication rate was then estimated by measuring the fluorescence of the strains during infection in *A. castellani* and U937 cells. Growth curves (**Figure 2A**) and growth rates (**Figure 2B**) indicate no growth deficit of OmpP1/FadL-variant bearing clinical isolate DA38626 compared to the environmental isolate DA38627 in *A. castellani* (average growth rates increase by 5.49%, corresponding to an decrease in doubling time of 3.7 minutes), yet a marked increase in growth rate in macrophages (average growth rate increase of 34.9%, corresponding to a decrease in doubling time of 40.6 minutes).

**Figure 2:**
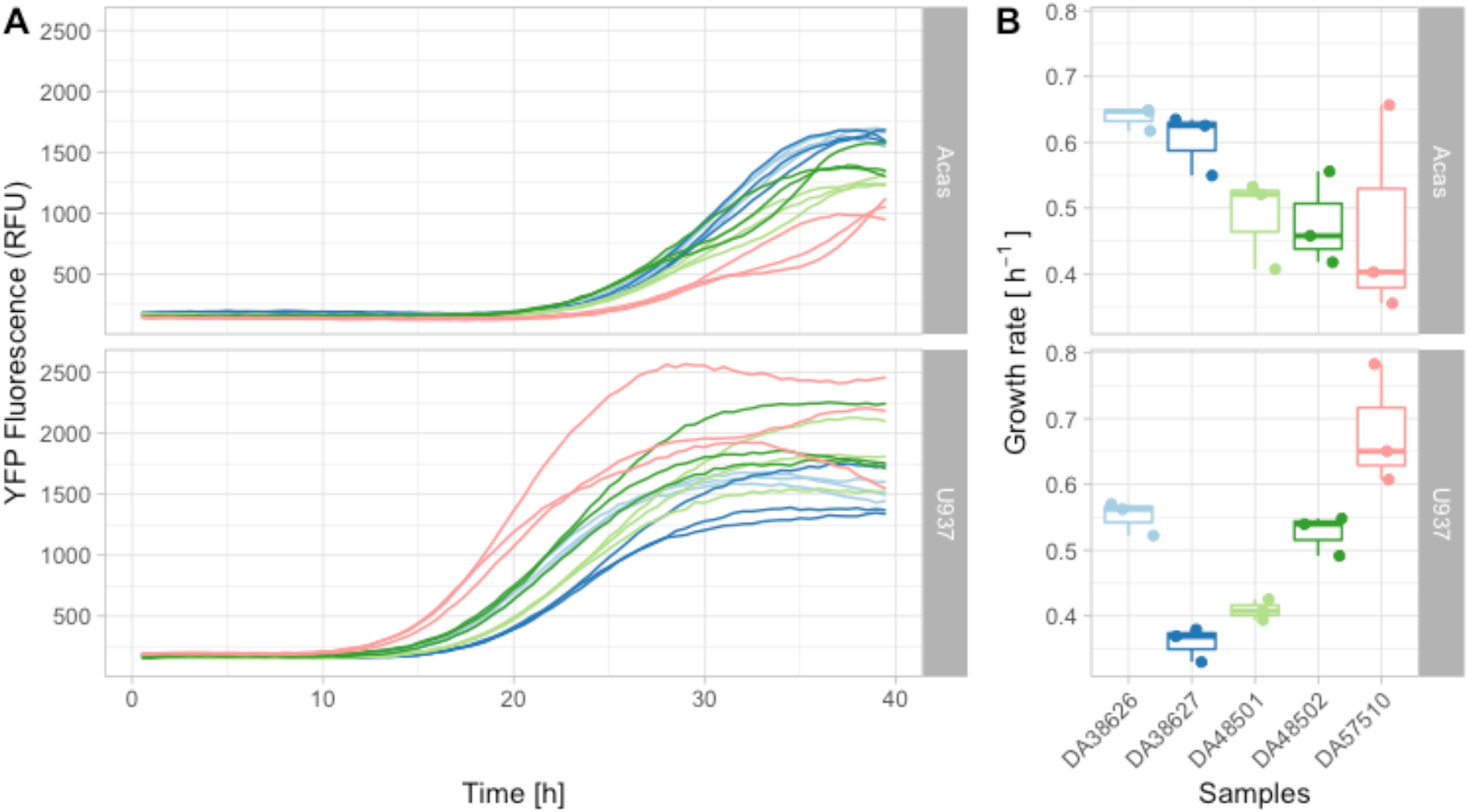
Intracellular replication of *L. pneumophila* in *A. castellanii* and U937 cells at an MOI of 50. Each dot or curve represents the average of three technical replicates, and each strain was measured with three separate biological replicates. DA38626 (clinical isolate, pale blue) carries the mutated OmpP1/FadL homolog (lpg0707), while DA38627 (environmental, dark blue) has the wild type gene. DA48501 (clinical isolate, pale green) carries a mutated EAL-containing protein (lpg0891), while DA48502 (environmental, dark green) has the wild type gene. DA57510 (red) is *L. pneumophila* str. Paris and serves as control. **A:** Growth, as measured by fluorescence (y-axis), over time (x-axis, in hours). **B:** Growth rates, as inferred from the growth curves by Growthcurver [39].

The EAL-variant bearing clinical isolate DA48501 compared to the environmental isolate DA48502 shows a moderate decrease in *A. castellani* cells (average growth rate increase of 2.05%, corresponding to an decrease in doubling time of 1.7 minutes) but, contrastingly, a more important decreased growth rate in macrophages (average growth rate decrease of 28.6%, corresponding to an increase in doubling time of 23 minutes). Interestingly, compared to all these isolates, the control *L. pneumophila* str. Paris grows slower in *A. castellanii* but faster in U397 cells.

## Discussion

By comparing a large number of *L. pneumophila* clinical isolates from outbreaks and sporadic cases to their environmental source we were able to identify nine genes, which acquired a mutation in more than one independent case, presumably during the infection of the patient. Of these, two were mutated three times. These genes showing signs of convergent evolution potentially represent adaptations of *L. pneumophila* to the human host.

We focused on SNPs, and found a high fraction (69%) of non-synonymous variants, indicating that selective pressure, and not only genetic drift, is at work [40, 41]. Another driver of genomic diversity is horizontal gene transfer (HGT). In *L. pneumophila* all three primary mechanisms of bacterial recombination (conjugation, transduction and transformation) have been described [21, 22, 42]. Recombination events are frequent and the exchange of large fragments (>200 kb) has been described [23]. In addition, within certain lineages 95% of SNPs arose due to recombination events [20, 25]. While stretches of mutations originating from recombination were found in our own samples, they did not account for any of the possible candidates of host adaptation. Although some of the isolated mutations included in our analysis might still come from recombinations, it would be from recombinations between very closely related strains, and does not affect the results of our analysis. Indeed, mutations resulting from recombinations may still represent adaptations.

Another complicating factor may have been the time between infection and environmental sampling [43], which could have led to the rise of variants in the environmental samples. By obtaining enough samples, and by removing comparisons from outbreaks in which the exact same mutation occurred in all comparisons, this effect is limited. Additionally, for both genes mutated three times, we always observed the same wild type allele in the environmental strain and a mutated one in the clinical strain.

The probability of obtaining as many as seven genes independently mutated without selection is low, but genes mutated twice appeared in all simulations. Since it is not possible to discriminate with high confidence, among the seven candidates, those that arose by chance alone from occurrences of convergent evolution, genes mutated twice are not discussed further. We focus on the two genes mutated three times, which occur only very rarely in simulations. These two candidates, encoding an outer membrane protein (OmpP1/FadL homolog; lpg0707) in which two out of three mutations encode an early stop codon and an EAL-domain contain (lpg0891), are discussed below. All mutations occurring in these two genes have presumably occurred either in-patient or slightly before in the same strain. Even mutations that occurred before infection in the direct ancestor of the strain that eventually infected the patient are relevant, since those mutations have a selective value and favor the infection.

The outer membrane of *Legionella* consists of an inner layer consisting of various phospholipids [44] and an outer layer of phospholipids and lipopolysaccharides (LPS) [45]. One study proposed the existence of around 250 proteins in the *L. pneumophila* outer membrane [46]. Most of their functions remain unclear, and this includes the outer membrane protein encoded by the gene lpg0707 identified in this study. This protein belongs to the OmpP1/FadL family, and was identified inside a recombination hotspot in the ST62 lineage [24]. The lipopolysaccharide (LPS) layer of Gram-negative bacteria is an effective barrier for the passage of hydrophobic molecules through the outer membrane. The FadL family of proteins is conserved throughout bacterial species and is required for the uptake of hydrophobic long-chain fatty acids via lateral diffusion through the outer membrane [47, 48]. In this study, two of the mutations cause a premature stop codon at AA position 26 and 245, potentially leading to the expression of a non-functional FadL. Based on a protein sequence alignment and the crystal structure of a *Pseudomonas aeruginosa* FadL homologue [47] these stop codons would disrupt the β-barrel that functions as a hydrophobic tunnel through which substrates pass (**Figure 3**). The third mutation does not cause a change in secondary structure, but replaces a glycine (small, non-polar) with a negatively charged aspartate at position 311, in the part located in the middle of the hydrophobic membrane. Although the exact effect of this mutation cannot be determined without further experimental evidence, it is not unreasonable to believe that it affects the function of the OmpP1/FadL protein in a significant way, possibly resulting in a loss-of-function mutation as the other two.

**Figure 3:**
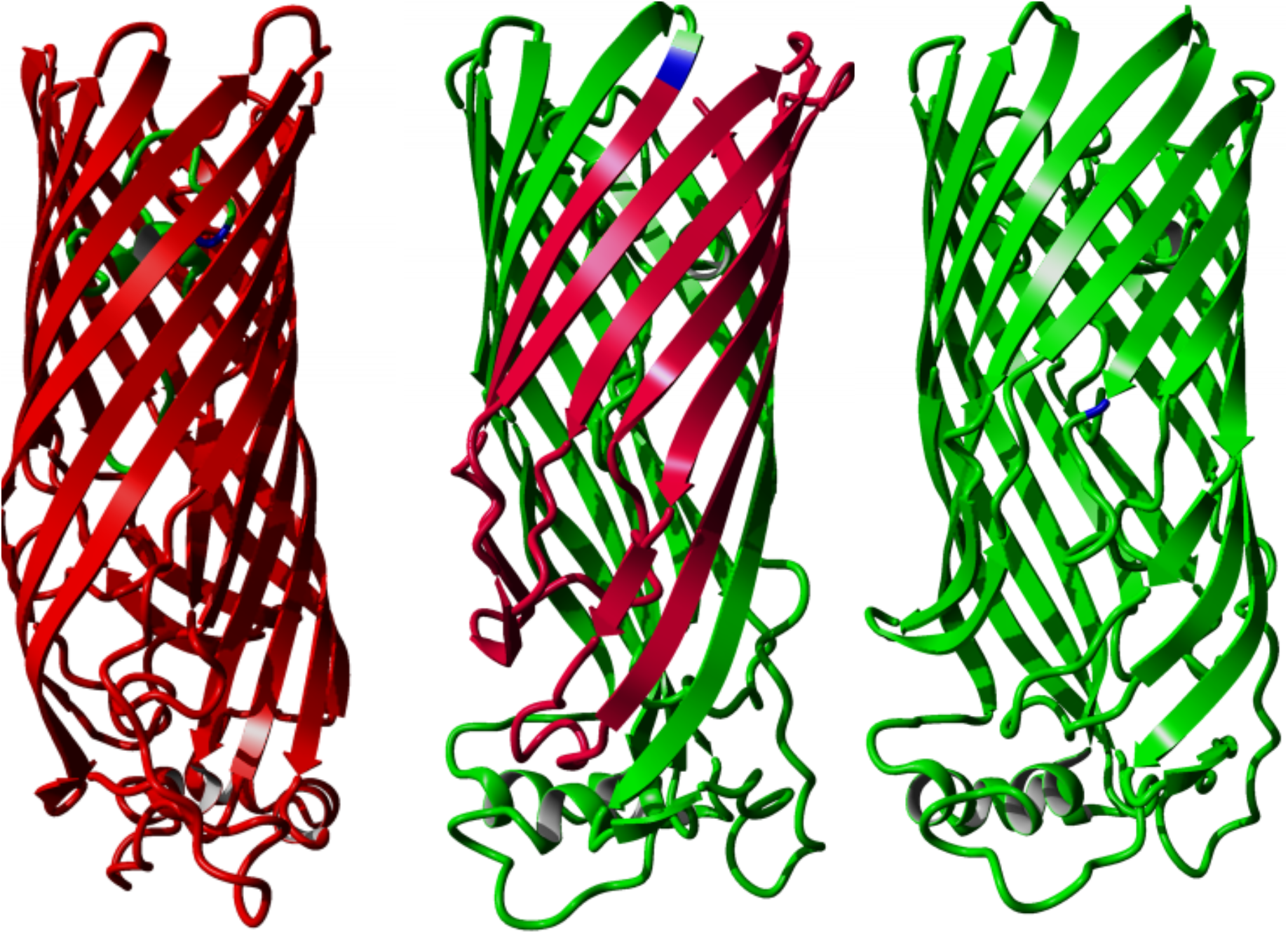
Crystal structure of *Pseudomonas aeruginosa* FadL homolog (PDB: 3DWO), with the corresponding location of the variants highlighted in blue (left: strain BrisbaneLP47, G26*, aligned to S28 in 3DWO; middle: strain PHHL01023034, W245*, aligned L300 in 3DWO; right: DA38626, G311D, aligned to T362 in 3DWO). Red indicates the part of the protein that would not be translated after the introduced stop codons.

Our growth assays show that the mutation in OmpP1/FadL confers a significantly faster growth (34.9%) in the human macrophage cell line U937, while providing very little difference in the natural host *A. castellanii*. This suggests that the mutant is better adapted to human macrophages, without a clear cost to its ability to infect and replicate in its natural amoebal host.

In a screening of *Salmonella paratyphi A* CMCC 50973 outer membrane protein, FadL was found to be strongly immunogenic and thus proposed as a candidate for a vaccine [49]. Tolllike receptors (TLRs) of the human innate immune system detect multiple pathogen-associated molecular patterns such as LPS, which is detected by TLR4 or TLR2 for *Legionella* [50, 51]. Therefore, modifications or disruptions to the LPS or the outer membrane could allow *L. pneumophila* to avoid early detection by the human host’s immune system, and allow it to establish itself inside the alveolar macrophages. However, since our experimental setup includes cells only, it appears unlikely that the mutation only allows *L. pneumophila* to effectively evade the immune system response. The loss of function in a channel-like protein like the OmpP1/FadL could reduce permeability to harmful substrates present in the macrophage cytosol but not in the amoebal one. For example, it has been shown that loss of the OmpP1/FadL homolog in *Haemophilus influenzae* reduces its uptake of bactericidal long-chain fatty acids (LCFA), increasing its resistance to the latter compound [52]. LCFA are, among others, produced by hydrolysis of the host membrane phospholipids. Further investigations of the function of this gene will shed light on its role in infecting the accidental human host.

EAL domain-containing proteins regulate cyclic diguanylate (c-di-GMP), a ubiquitous second messenger in bacteria. C-di-GMP mainly modulates pathways involved in lifecycle transitions, e.g. the transition from the motile to the sessile state, the transition from pathogenic to environmental lifestyles, or from planktonic infections to biofilm infections [53–57]. Cyclic-di-GMP is produced from GTP by diguanylate cyclases (DGCs), and is degraded via phosphodiesterases (PDEs). The catalytic site of DGCs is identified as a GGDEF domain, while PDE activity involves either an EAL or the HD-GYP domain [58–60]. *L. pneumophila* encodes 22 to 24 GGDEF/EAL proteins, and most are highly conserved in *L. pneumophila* strains, suggesting an important role of GGDEF/EAL domain proteins for regulation of the *L. pneumophila* life cycle [57, 61].

Upon infection, *Legionella* uses its Dot/Icm type 4 secretion system (T4SS) to secrete over 300 effector molecules to establish a vacuole called the *Legionella*-containing vacuole (LCV), in which it may replicate. The LCV allows *L. pneumophila* to replicate until nutrient deficiencies lead to a reprogramming genetic expression. This includes the expression of virulence factor-encoding genes, and the upregulation of genes containing GGDEF/EAL domains [62–64]. The specific protein encoded by lpg0891, mutated independently three times in this study, harbors a DGC domain and PDE domain, both functional [65]. Two out of 3 mutations occurred within an EAL domain (**Figure 4**), but none apparently resulted in a loss-of-function mutation. All three mutations replace a non-polar amino-acid by a polar one. In strain Philadelphia, the protein was shown to be expressed only in the post-exponential phase, i.e. at the end of the intracellular replication, when *Legionella* is virulent and motile [66]. In the same strain, knock-out mutants for this protein are still able – albeit to a slightly lesser extent – to grow in BYE broth, and to infect both amoebae and macrophages, to translocate and to evade lysosomes [65], but experiments in strain Lens have demonstrated that the homolog protein encoded by lpl0922 is crucial for setting up efficient intracellular replication, both in amoebae and macrophages [67]. It has also been shown that the homolog protein in strain Paris (encoded by gene lpp0952) plays an important role in the regulation of flagellar activity and of motility, and is regulated by the flagellar regulator FleQ, presumably via FliA [68].

**Figure 4:**
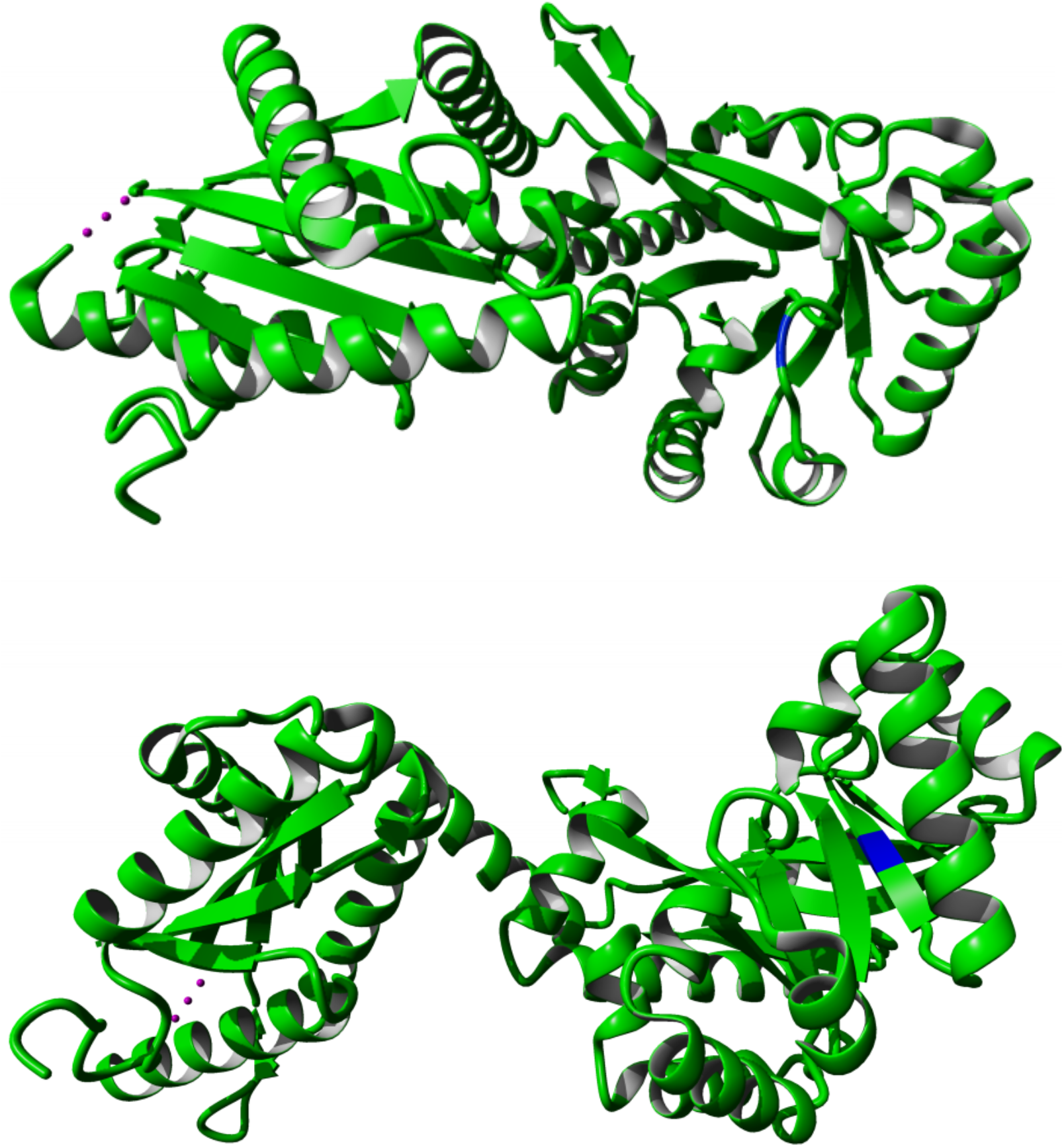
Crystal structure of *Pseudomonas aeruginosa* MorA homologue (PDB: 4RNH), with variants highlighted in blue. Top: G662D mutation in strain PHHL01023035, aligned to G1315. Bottom: I711T in strain DA48501, aligned to I1365. The G157R mutation in strain PHH072360604 does not align to the crystallized part of MorA and is thus not shown here.

The EAL-domain containing protein mutant studied here displays a similar growth rate as the wild type in the amoebal host, but an important reduction in growth rate (28.6%) when infecting human macrophage cell lines. This surprising result suggests that the selective pressures that favored the mutant allele in the three independent infections happened outside of the cell. This is however consistent with the fact that many of the functions related to EAL domains in *Legionella* have a role in regulating mobility outside the cell, e.g. through flagellar regulation. However, the exact role of this gene remains elusive, and further functional characterization is required to shed light on the mechanisms by which the mutated allele provides a better adaptation to the human host.

A recent extensive study systematically explored the importance of *L. pneumophila* genes for growth in four amoebal hosts and one human histiocytic lymphoma cell line: *Acanthamoeba castellanii, Acanthamoeba polyphaga, Hartmannella vermiformis, Naegleria gruberi*, and U937 [28]. This study highlighted that different hosts impose different evolutionary constraints on different hosts, with gene or gene variants being beneficial in some hosts and detrimental in others. None of the genes identified in our study correlated to any of the genes highlighted in the study mentioned, but this study did not consider the full human immune system and lpg0891 and lpg0707 may still play an important role in infecting human hosts. It also emphasizes the possibility that a single mutation may carry a large selective value in presence of a specific host, especially one that is not often met.

The exact population size of *Legionella* inside the human host is unknown. A study using guinea pig models found that a reproducible infection was observed for an estimated dose of ≥5 CFU retained in the lungs [69, 70]. However, the inhaled dose compared to the deposited dose is unknown, as is the composition of the *Legionella* population inside human lungs. One study on ten patients found no to very low within-host diversity [71], an hypothesis confirmed by more studies [72, 73]. Other studies, however, found the opposite [38, 74], and it is thus likely that within-host diversity depends on the diversity of *L. pneumophila* in environmental sources, variations in infectious dose, and the duration of infection prior to sampling [71]. While antibiotic resistance does not appear to be a current general concern in the environment, antibiotic resistance to fluoroquinolones has been shown to occur within patients [75].

In summary, we identified at least two potential candidate genes for in-patient, human-specific adaptation in *L. pneumophila*. This included a regulatory protein able to both synthesize and degrade cyclic-di-GMP, which is involved in the virulence and motility of *Legionella*. We also identified signs of convergent evolution in a OmpP1/FadL outer membrane protein, in which mutations led to several premature stop codons, and provided *L. pneumophila* with a large increase in growth rates in human macrophages. The repeated detection of mutation in the same genes, probably occurring during the short time (2-14 days) of the onset of the disease in patients, is consistent with these mutations being very beneficial to *L. pneumophila* when infecting human hosts. This suggests that *L. pneumophila* is not as well adapted to the human host than to the amoebal host.

## Material and methods

### Bacterial samples

Clinical (grown from sputum samples) and environmental samples were shipped to Uppsala by the Public Health Agency of Sweden (48 samples), the French National Reference Center of *Legionella* (96 samples), and the Spanish Laboratorio de *Legionella* (27 samples) on Buffered Charcoal Yeast Extract (BCYE) agar plates. Bacteria were collected directly from these plates, to minimize the number of generations and the accumulation of non-relevant mutations prior to sequencing.

### DNA isolation and sequencing

DNA was extracted from 154 clinical and environmental samples using the MasterPure™ DNA Purification Kit (Epicentre), following the manufacturer’s protocol. The quantity and quality of extracted DNA was assessed by NanoDrop (ThermoFisher) and gel electrophoresis on 1% agarose gel. Extracted DNA was prepared and sequenced using MiSeq v3 (Illumina), using 300bp paired-end sequencing at the National Genomics Infrastructure (NGI) Sweden, SciLifeLab, Stockholm, Sweden.

A literature study identified 66 published *L. pneumophila* samples for which clinical and potential environmental samples could be identified [20, 31–34, 36, 73, 76, 77] (**Supplementary Table 1**).

### Assembly

Read quality was controlled using FastQC [78] and MultiQC [79]. Reads were trimmed with SeqPrep [80], and de-novo assembled using SPAdes 3.9.1 [81]. Contigs of short length (<500 bp) and low coverage (<10X) were removed and remaining contigs were annotated using prokka 1.12-beta [82].

### Variant calling

Genetic variants (SNPs and INDELS) in clinical samples were called with RedDog [83], using the corresponding assembled environmental samples as reference. Samples with >20 SNPs were removed from the analysis, as in these, the time of divergence between the environmental strain and the patient strain is presumably too long and the high and most SNPs are likely to have occurred outside of the patient. As a control all samples were subjected to self-to-self variant calling, in which variants were called with the reads for each sample against their assembly. Samples with >10 SNPs called against themselves were removed from the analysis. The annotated genomes and RedDog variant positions were used to confirm if mutations were synonymous or non-synonymous.

### Ortholog clustering

All proteomes from the SPAdes assemblies and from 5 reference genomes (Alcoy, NC_014125; Corby, NC_009494.2; Philadelphia, NC_002942.5; Lens, NC_006369; Lorraine, NC_018139.1) were clustered into ortholog protein families using OrthoMCL [84]. Variants were then matched to their respective proteome to identify the consequences of the mutations, and to which protein family they belonged. Annotation of the protein families in which multiple variants were found between independent samples was done by searching for the protein accession numbers in the first reference genome and using each region as a query for a BLAST search.

### Simulation of non-selective environment

To estimate the distribution of mutations per gene to expect in a non-selective environment, 123 genes (which corresponds to the observed number of SNPs in this study) were sampled randomly, with replacement, from the 3033 genes harbored by *L. pneumophila* Paris. The number of genes sampled once or more times was recorded. The sampling was repeated 1000 times. The probability that the observed number of genes mutated twice and three times, respectively, would be greater than in a non-selective environment was assessed by comparing the actual number of genes to distribution obtained from the 1000 simulations, using one-sided Mann-Whitney tests, wilcox.test in the stats package of R [85]. The results were visualized with ggplot2 [86].

### Phylogenomics

Phylogenetic tree was generated based on a core genome MultiLocus Sequence Typing (cgMLST) scheme via the ChewBBaca pipeline [87]. The already available *L. pneumophila* Philadelphia (NC_002942.5) strain training file was used [88]. Loci present in 95% of genomes were included. The nucleotide sequences were aligned using MAFFT-LINSI [89]. A phylogenetic tree was produced using FastTree [90] with the Jukes-Cantor substitution model.

### Functional annotation

An amino acid sequence search in the RCSB Protein Data Bank [91] was used to identify homologous structures, protein sequences were then aligned using MAFFT-LINSI and the structural effect of the mutations were predicted via NetSurfP-2.0 [92]. Mutations were visualized in crystal structures of a *Pseudomonas aeruginosa* homologs (PDB accessions: 3DWO for the OmpP1/FadL homolog and 4RNH for the EAL-containing protein) via YASARA View [93].

### Bacterial strains, cell culture and media

*Legionella* strains used were cultured in charcoal yeast extract (CYE) (1% ACES, 1% yeast extract, 0.2% charcoal, 1.5% agar, 0.025% Iron (III) pyrophosphate, 0.04% L-cysteine, pH 6.9) plates or ACES yeast extract (AYE) (1% ACES, 1% yeast extract, 0.025% Iron (III) pyrophosphate, 0.04% L-cysteine, pH 6.9) broth at 37°C, unless otherwise stated.

To measure fluorescence as a proxy to bacterial replication, selected strains were transformed with a YFP-carrying plasmid, with the fluorescent gene under the IPTG-induced lacI promoter – pXDC101 (kindly given by Dr. Elisabeth Kay, University of Lyon). Transformation was done via electroporation to cells that were made electrocompetent by washing three times with 10% glycerol at 4°C. Positive mutants were henceforth always grown in media supplemented with 8 μg ml^-1^ Chloramphenicol (Cam) and IPTG (1 mM in solid, and 0.5 mM in liquid medium).

*Acanthamoeba castellanii* (ATCC 30010) was cultured in Peptone Yeast Glucose (PYG) medium (2% bacto proteose peptone 2, 0.1% yeast extract, 0.1% sodium citrate dihydrate, 0.4 mM CaCl_2_, 4 mM MgSO_4_ · 7H_2_O, 2.5 mM Na_2_HPO_4_ · 7H_2_O, 2.5 mM KH_2_PO_4_, 0.05 mM Fe(NH_4_)_2_(SO_4_)_2_ · 6H_2_O; 100 mM glucose, pH 6.5) in flasks at 30°C. For infections *A. castellanii* was washed and resuspended in LoFlo medium (ForMedium, Norfolk, UK), which does not support *Legionella* growth.

Human monocyte-like U937 cells (ATCC) were maintained in RPMI1640+GlutaMAX™ (Gibco) supplemented with 10% heat inactivated foetal bovine serum (FBS) (Gibco) and 1% Penicillin-Streptomycin (PS) (Gibco), in a 37°C incubator with 5% CO_2_. Before infection cells were harvested, viability and density were assessed using 0.4% Trypan blue solution (Gibco) and an automated cell counter (Countess™ FL, ThermoFisher). The cells were then centrifuged at 200 x*g* for 5 min, and resuspended in growth medium with 50 ng ml^-1^ of phorbol 12-myristate 13-acetate (PMA) for 48h to induce differentiation into macrophage-like cells. The medium was then replaced with fresh medium and cells incubated for a further 48h. On the day of infection the medium was changed to infection medium: RPMI 1640 without phenol red (Gibco) supplemented with 10% heat inactivated FBS and 1% GlutaMAX™ (Gibco), this medium also does not support *Legionella* growth.

### Extracellular growth assays

*L. pneumophila* strains were grown on CYE plates at 37°C for 48h, resuspended in AYE, and diluted to an absorbance at OD_600_ of 0.1 (ca. 2×10^8^ c.f.u.). In a 96-well plate, 200 μl (ca. 4×10^7^ c.f.u.) of each cell culture was aliquoted per well (5 replicates each). Growth rate was tracked by measuring OD_600_ and YFP fluorescence (Excitation: 508 nm, Emission: 555 nm) every 30 minutes for 72h at 37°C, and 180 rpm shaking, using a Tecan Spark™.

### Infection assays

To assess the intracellular replication of the different *Legionella* strains, *A. castellanii* and U937-derived macrophages were challenged at different MOIs.

On the day of infection *A. castellanii* were harvested, washed, and resuspended in LoFlo. Then cell viability and density was evaluated as described above; 1×10^5^ cells were seeded per well in a 96-well plate, and the plate was incubated for 1h at 30°C, before adding bacteria.

Two hours prior to infection, each well, with previously seeded and differentiated U937 cells (1×10^5^ cells per well in a 96-well plate), was washed with DPBS (Gibco), and infection medium was added, the plated was incubated at 37°C with 5% CO_2_ for 1h.

*Legionella* strains used for infection were grown on CYE plates for 72h, resuspended in the respective infection medium for each host, OD_600_ was measured, and cultures were serial diluted to obtain desired cell density. Bacterial dilutions were then added to host cells, in a ratio of 10 bacteria: 1 host cell. Each strain was tested in 5 replicates.

To track the increase of *L. pneumophila* during infection plates were incubated on a Tecan Spark™ at 30°C, or 37°C with 5% CO_2_, and YFP fluorescence was recorded every 30 minutes for 72h.

### Growth rate analysis

Growth rates were calculated by importing the fluorescence and OD data into R [94] and analyzed with Growthcurver v0.3.1 with default settings [39].

## Author statements

### Authors and contributors

L.G. conceptualized the study, acquired funding, and supervised the project. C.P., M.M., E.H, D.K., S.J., and C.G. isolated and characterized bacterial isolates. D.L. obtained and curated the data, performed the bioinformatic analysis, visualized and analyzed it, with contributions from S.M. and L.G. A.B.M. designed, performed and analyzed the experimental part. D.L. wrote the original draft. All authors contributed to reviewing and editing the manuscript. All authors approve the final version of the manuscript.

### Conflicts of interest

The authors declare that there are no conflicts of interest.

### Funding information

This project is funded by grants to L.G. from the Swedish Research Board (VR, grant 2017-03709) and the Carl Tryggers Foundation (grant CTS 15:184), and by a scholarship from the Japan Society for the Promotion of Science to D.L. Sequencing was funded by the SciLifeLab program “Swedish Genomes and Biodiversity” (2015). Funders had no role in designing the study or interpreting the results.

## Acknowledgments

The authors would like to thank Matilda Morin, Elisabeth Hallin and Daniela Klingenberg (Public Health Agency of Sweden), as well as Carmen Pelaz (Legionella Reference Laboratory in Spain) for providing strains from their collection; Ulrika Lustig for her patient technical assistance; and Elisabeth Kay for providing the pXDC101 plasmid. The authors would also like to acknowledge support from Science for Life Laboratory, the National Genomics Infrastructure, NGI, and Uppmax for providing assistance in DNA sequencing and computational infrastructure.

## Supplementary material

Supplementary Table 1

Isolates sequenced in this study. Separate Excel file.

Supplementary Table 2

Comparison clinical-environmental isolates analyzed in this study. Separate Excel file.

## Notes

https://www.ncbi.nlm.nih.gov/bioproject/PRJEB52976

### Competing Interest Statement

The authors have declared no competing interest.

